# Estimating intra-subject and inter-subject oxygen consumption in outdoor human gait using multiple neural network approaches

**DOI:** 10.1101/2024.04.25.591094

**Authors:** Philipp Müller, Khoa Pham-Dinh, Huy Trinh, Anton Rauhameri, Neil J. Cronin

**Affiliations:** Faculty of Information Technology and Communication Sciences, Tampere University, 33720 Tampere, Finland; Faculty of Medicine and Health Sciences, Tampere University, 33720 Tampere, Finland; Faculty of Sport and Health Sciences, University of Jyväskylä, 40014 Jyväskylä, Finland

## Abstract

Oxygen consumption 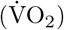 is an important parameter for exercise test, such as walking and running, that can be measured using portable spirometers or metabolic analyzers. However, these devices are not feasible for regular use by consumers as they intervene with the user’s physical integrity, and are expensive and difficult to operate. To circumvent these drawbacks, indirect estimation of 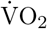 using neural networks combined with motion parameters and heart rate measurements collected with consumer-grade sensors has been shown to yield reasonably accurate 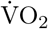 for intra-subject estimation. However, estimating 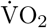 with neural networks trained with data from other individuals than the user, known as inter-subject estimation, remains an open problem. In this paper, five types of neural network were tested in various configurations for inter-subject 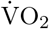 estimation. To analyse predictive performance, data from 16 participants walking and running at speeds between 1.0 m/s and 3.3 m/s were used. The most promising approach was XceptionNet, which in most configurations even yielded a lower average estimation error than the LSTM neural network from an earlier study for intra-subject estimation. This suggests that XceptionNet could be embedded in portable devices for real-time estimation of oxygen consumption during walking and running.

## Introduction

Oxygen consumption (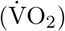), also known as oxygen uptake, is frequently used to measure walking and running economy since the exchange of oxygen and carbon dioxide is highly correlated to energy metabolism. For unconstrained walking or running 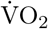 can be measured directly by metabolic analyzers or portable spirometers, but these devices are inconvenient to use regularly, often require trained personnel for operation, and are expensive.

Therefore, research on indirect estimation of 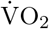 from observations of surrogate variables has received a lot of attention over the last two decades. Indirect estimation has benefitted from recent advances in machine learning techniques and development of consumer-grade, small, wearable sensors. The most common approach for 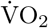 estimation is based on heart rate (HR) measurements [1]. For example, several commercial products, such as the Suunto personal HR monitoring system, use HR data for estimating 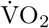 and energy expenditure [2]. Because HR is affected by age, sex, fitness level, exercise modality, environmental conditions, and day-to-day variability [3 from Pavel], HR index (HRI) is frequently used instead of HR [1, 3]. HRI is obtained by dividing the HR measurement by an individual’s resting HR, which has the potential to remove the need for individual calibration [3]. In [2]additional features such as R-wave-to-R-wave (R-R) heartbeat intervals, R-R-based respiration rate, and on-and-off 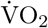 dynamics at various exercise conditions were used for 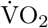 estimation. However, authors of the study acknowledged the limitations in the estimation accuracy when including individual maximal 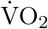 and HR values. Several studies have used linear regression models to estimate 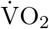 [1–4], which worked well for moderate intensity exercises. However, for very low and very high intensity exercises the relationship between 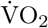 and HR is significantly nonlinear, resulting in poor 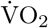 estimates.

Other factors that can affect the relationship between 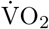 and HR include altitude, exercise duration, hydration status, medication, state of training, and time of day [2 from pavel]. To account for these factors, in cycling breathing frequency, mechanical power, and pedaling cadence, which can be measured directly from cycling ergometers, can be included for 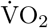 estimation [5–7]. For walking or running in unconstrained outdoor environments, breathing frequency, cadence, speed, and speed variation calculated by wearable devices can be used as input features [4].

In [8] we computed motion parameters, namely step-wise average speed, peak-to-peak speed difference, step duration, and peak-to-peak difference in vertical movement from measurements of an inertial navigation system combined with a Global Positioning System (INS/GPS) device. The wearable INS/GPS device measured acceleration, velocity, angular velocity and orientation of the upper body (for details the reader is referred to [4]. The four motion parameters were used together with HR as input features for estimation of 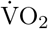 during walking and running by a long short-term memory (LSTM) neural network. The results suggest that LSTM neural networks are able to accurately estimate oxygen consumption; the achieved accuracy was 2.49 ml×min^−1^×kg^−1^ (95% limits of agreement). For comparison, in [10] 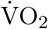 during walking and other daily activities were estimated by random forest regression using breathing frequency, HR, hip acceleration, minute ventilation, and walking cadence; the achieved accuracy was 6.17 ml×min^−1^×kg^−1^ (95% limits of agreement).

One limitation of the study in [8] was that data for training the LSTM neural networks and for evaluating its performance came from the same individual (intra-subject estimation). However, estimating oxygen consumption for an individual by a model trained with data from the same individual is time-consuming and not always feasible. Ideally, the model would already be trained beforehand, using training data from other individuals, and used immediately for 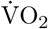 estimation. This is referred to as inter-subject estimation. Furthermore, in [8] only light and moderate intensity exercises were covered.

Therefore, in this paper a wide selection of neural network models are studied for estimating inter-subject oxygen consumption across a range of walking and running speeds (1.0 m/s to 3.3 m/s) based on measurements of motion parameters and heart rate. The contributions of our paper are three-fold. First, we demonstrate that by using an early exit strategy and optimizing hyperparameters the accuracy of the LSTM model from [8] can be significantly improved (average estimation error was reduced by approximately 82%). Second, we show that with more sophisticated neural network structures 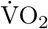 estimates for inter-subject estimations can be obtained that are more accurate than the intra-subject estimations yielded by the LSTM model from [8]. Finally, a more detailed correlation analysis between the neural networks’ input features (motion parameters and HR) and output feature 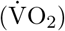 than in [8] is provided, which yields insights into why neural networks are able to yield accurate 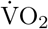 estimates.

## Materials and methods

### Experimental data

Sixteen healthy participants between 18 and 35 years of age (age 27.5 ±3.5 yrs, height 175.3 ±8.4 cm, body mass 71.8 ±12.9 kg, body mass index 23.3 ±3.4 kg/m^2^, eight females) participated in the field tests on a level outdoor track. Ten of the participants were recreational runners, meaning that they ran at least twice per week during summer and performed other endurance sports during winter. Their statistics were: age 28.1 ±3.7 yrs, height 177.6 ±6.4 cm, body mass 70.8 ±11.4 kg, body mass index 22.3 ±2.7 kg/m^2^, five females. For the remaining six participants age 26.5 ±3.1 yrs, height 171.5 ±10.59 cm, body mass 73.3 ±16.1 kg, body mass index 24.8 ±4.2 kg/m^2^, three females. These participants ran at most twice per month. The Ethics Committee of the University of Jyväskylä approved the study. Participants were recruited between 5 May 2018 and 31 July 2018. All participants were informed about the content and purpose of the testing procedure, and provided written informed consent, witnessed by one researcher. The research was conducted in accordance with the World Medical Association Declaration of Helsinki [11].

Each participant was equipped with a datalogger, a portable spirometer, and a chest strap for measuring the heart rate. The datalogger was assembled on a Raspberry Pi 3 model B running Raspbian operating system, connecting with a high quality Vectornav VN-200 (Vectornav Technologies, United States) GPS-aided inertial navigation system (INS/GPS), a GPS antenna, and a battery. The IMU incorporates an accelerometer, a gyroscope, a magnetometer and a barometric pressure sensor (details can be found in [4]). The device measures 150 x 75 x 48 mm and was carried on the participant’s upper back in an orienteering battery vest. Oxygen consumption and other breathing variables were measured during the walking, running and rest periods with a Jaeger Oxycon Mobile portable breath gas analyser (Viasys Healthcare GmbH, Germany). The setup consists of a desktop and a portable setup, with the latter including a sensorbox, a data exchange unit, and a mask to which a digital volume transducer and a gas tube were connected. Both sensorbox and data exchange unit were carried on the participant’s upper back with a special vest so that the units were located on either side of the datalogger. Heart rate was measured using a Polar V800 heart rate monitor and an H10 strap with integrated heart rate sensor (Polar Electro Oy, Finland).

Participants were asked to rest for five minutes at the beginning of the measurement session to obtain oxygen consumption at rest, which enabled studying the effect of exercise on a participant’s oxygen consumption. After that, participants were asked to walk or run along a 200 meter long track on the main straight of the level outdoor track at various speeds. Walking speeds were 1.0 m/s, 1.3 m/s, and 1.5 m/s; running speeds were 2.2 m/s, 2.5 m/s, 2.8 m/s, 3.1 m/s and 3.3 m/s. Subject 3, in addition, also ran at 3.6 m/s. Each participant started with walking at 1.0 m/s. The order of the remaining seven speeds were randomized for each participant individually by a browser-based randomizer (http://www.random.org/lists). Speed was controlled by LED modules spaced at one meter along the track, which enabled control of speed with an accuracy of 0.1 m/s. Participants were asked to follow the lights while walking/running for five minutes for each speed. After each walking/running speed, participants stopped for a few seconds and then returned to the starting point to sit still for five minutes, allowing heart rate and oxygen consumption to return to resting levels.

The Oxycon Mobile spirometer measured every five seconds 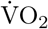 and respiratory frequency using breath-by-breath methods. To ensure accurate measurements, the spirometer was re-calibrated at the start of each measurement session.

In line with [8], oxygen consumption measurements were smoothed by applying a Savitzky-Golay filter [12] with polynomial order and window length set to 3 and 1 respectively three times. Due to unusually noisy data, for subject 7 also polynomial order set to 9 was tested and used in inter-subject estimations. Heart rate was recorded continuously (beat-by-beat) during the test at a sampling rate of 1 Hz. Data were smoothed and interpolated after the tests. Smoothing was done by a moving average with window length 3. For subjects 4 and 9 the window length was increased to 5 due to the exceptionally noisy heart rate data. The INS/GPS datalogger recorded acceleration, velocity, angular velocity and orientation at 400 Hz and saved them to a memory card through a wired connection, preventing any data loss. Accuracy levels of speed and speed difference were approximately 0.05 m/s; accuracies of computed vertical oscillation and step duration were about 1 cm and 10 ms respectively (for more details refer to [4]).

After the measurement campaign it was noticed that for subject 1 heart rate data was only partly available. Thus, only parts for which heart rate data as well as data from the INS/GPS datalogger and the portable spirometer were available were used for analysis. Similarly, the dataset for subject 2 lacked spirometer data for approximately 90 s, thus all data from this period was removed during data preprocessing. Subject 10 terminated the measurement campaign after the fifth walking/running cycle. Due to the limited number of participants, data from these five cycles were, nevertheless, included in the analysis.

### Dataset preparation

After data collection, measurements of spirometer, INS/GPS datalogger, and heart rate device were synchronized in time. Synchronization of spirometer and INS/GPS datalogger data was based on the internal clocks of both devices. For oxygen and heart rate time series data their cross-correlation was calculated and the highest peak was used as offset estimate for synchronization.

Step segmentation described in [4] was applied to motion data from the INS/GPS datalogger and used to compute walking/running metrics commonly used in gait analysis on a step-by-step basis (see [4] for details). Accelerations and velocities were computed in the anatomical frame. In the anatomical frame the x-axis is pointing into the direction of progression (anterior direction) and the z-axis is pointing upwards, parallel to the field of gravity. The y-axis is perpendicular to x- and z-axes and completes a right-handed coordinate system. Oxygen consumption and heart rate measurements were resampled to match the step-by-step frequency of walking/running metrics.

In [8] feature engineering was used to identify input features derived from the INS/GPS data that are strongly correlated with oxygen consumption, the so-called target feature, but only weakly correlated with other features. Based on the results in [8], for each step the following five input features were computed to be used in training neural networks for oxygen consumption estimation:

- Speed: arithmetic mean of the velocity (=step length/duration of step) over one step, measured in m/s
- Speed change: peak-to-peak difference in speed during one step, measured in m/s
- Step duration: measured in s
- Vertical oscillation: peak-to-peak difference in vertical movement, measured in m
- Heart rate: measured in bpm

Sequences of steps were used as inputs for the network training. The experiments for [8] indicated that input sequences of 50 steps yielded satisfactory estimates of the time-dependent decay between oxygen consumption (target feature) and past input values, thus the same length was used in this paper as starting point. The input sequences were normalized to values between 0 and 1 to enable better fit and prevent divergence in the network training.

### LSTM network architecture for intra-subject estimations

In [8] a many-to-one long short-term memory (LSTM, [17]) model was developed and tested successfully for intra-subject oxygen consumption (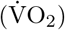) estimation. The network structure was simple and consisted of a sequence input layer, a LSTM layer with 150 hidden units, one dense layer, and an output layer. For training the network the *Adam* optimizer was used; the learning rate was set to 0.005; and training was run over 8 000 epochs. The motivation for using such a large number of epochs was that only a small dataset was available (subjects only walked/ran four times three minutes at four different speeds). Since a more extensive dataset has been collected for this paper, the *LSTM model* from [8] was trained here for 1 000 epochs only.

For [8] the aim was to demonstrate that LSTM networks can successfully estimate oxygen consumption, rather than finding the most accurate network modelFor this paper several alternative network structures were tested. First, twelve simple modifications of the *LSTM model* were studied. The most promising modification was an Early Exit Neural Network (e.g. [9]) that used 100 instead of 150 hidden units in the LSTM layer. The initial learning rate of 0.005 was decreased by factor 0.2 if the validation loss did not improve over the last 25 epochs. The minimum learning rate was set to 10^−6^. Furthermore, in order to avoid overfitting, training was terminated if over the last 20 epochs no sequence of three epochs with decreasing difference between training and validation loss was observed. Hereafter, this model is referred to as *Modified LSTM model*.

The *LSTM model* [8] was compared with the *Modified LSTM model* by analysing their performances for intra-subject oxygen consumption estimation. Data from each of the 16 subjects was divided into training (70% of all samples), validation (15%), and test datasets (15%) randomly. After that for each subject both the *LSTM model* and the *Modified LSTM model* were trained. While the *LSTM model* was always trained for 1 000 epochs, *Modified LSTM model* was trained for 1 000 epochs or until the termination rule described in Subsection was fulfilled. The 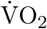 estimation capabilities of the both network structures were evaluated using the corresponding test datasets. This process was repeated for each subject five times (5-fold cross-validation) to assess how well the model generalises to different parts of the same participant’s dataset.

### Network architecture for inter-subject estimations

For inter-subject estimations the simple neural networks used for intra-subject estimations are insufficient, since the physiological and biomechanical features of gait differ between individuals. Therefore, in this paper a model architecture with a mandatory regression network and an optional feature network was used for a selection of more sophisticated neural network types. This architecture allowed the model to process both temporal and categorical input data. Temporal data included the same features as for the networks meant for intra-subject estimation, namely speed, speed change, step duration, vertical oscillation, and heart rate. Categorical data consisted of the following four individual features of test subjects:

- age: set to 0 for age up to 25, to 1 for age from 26 to 29, and to 2 for age of 30 or higher
- body mass index (BMI): set to 0 for BMI under 22, to 1 for BMI from 22 to 25, and to 2 for BMI over 25
- sex: set to 1 for male and 0 for female
- fitness level: set to 1 for recreational runners and 0 otherwise

The limits for categorical variables age and BMI were chosen based on the age and BMI distribution amongst the participants, to ensure approximately equal group sizes. Use of categorical data was optional, and each network type was tested with and without participant-specific features as input parameters. Fig 1 illustrates the two-headed model architecture. In case participant-specific features were ignored, only the branch of the regression network was trained and used to estimate oxygen consumption.

**Fig 1.**
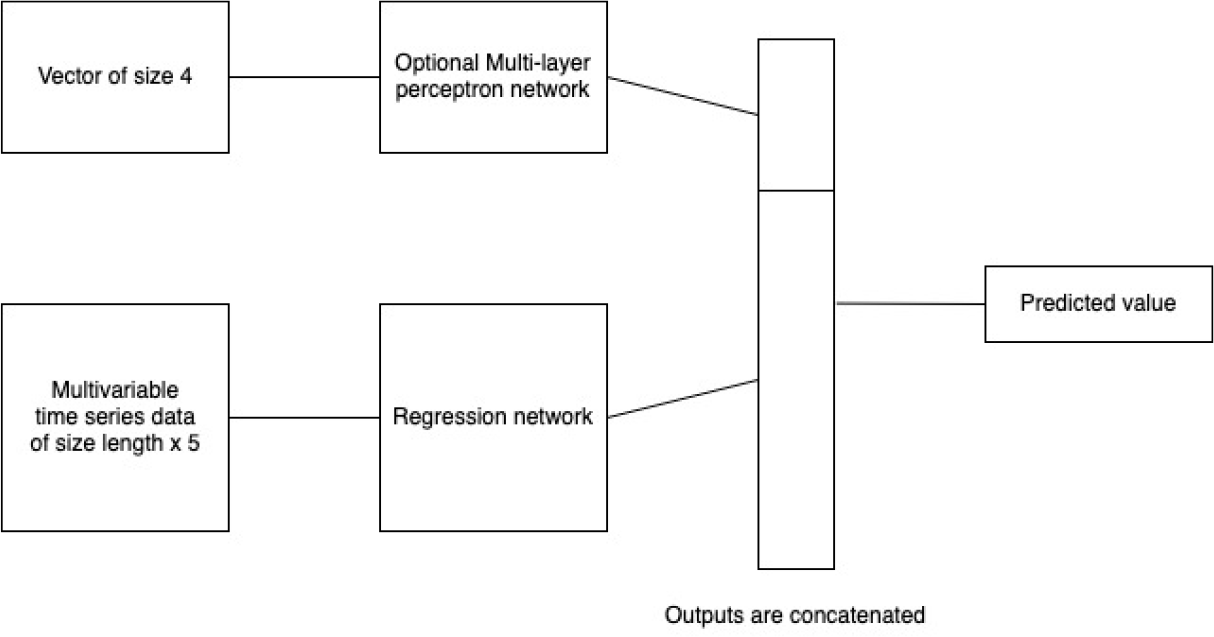
Network architecture used for inter-subject. 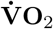 **estimation**. Upper, optional branch used four categorical variables age, body mass index, sex and whether or not the individual was a recreational runner as input data. Lower branch used temporal data for speed, speed change, step duration, vertical oscillation, and heart rate as input. Estimated value was 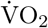.

Each network type was furthermore trained in various configurations. Due to the relatively small size of the dataset 100 epochs proved to be sufficient for achieving convergence in the network training phase. For all networks the AdamW optimizer [13] and the cosine learning rate scheduler with an initial learning rate of 10^−3^ and a final learning rate of 10^−5^ were used. Batch size was set to 64.

For evaluating the network types and configurations for inter-subject 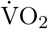 estimation, the following procedure was used. For each of the sixteen participants data from the remaining fifteen participants were used to train (data from thirteen participants) and validate (data from two participants) the networks and afterwards 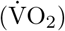 for the sixteenth subject was estimated with the trained models (leave-one-out cross-validation). This process was repeated five times for each of the sixteen participants (5-fold cross-validation). In the end the average root-mean square error (RMSE) and its corresponding standard deviation over all sixteen participants and five repetitions per participants were calculated.

#### Multi-layer perceptron for processing participant-specific features

The feature head is a multi-layer perceptron (MLP) with an output of two neurons. The network is described in Fig 2. A four-dimensional vector containing participant-specific features was used as input vector. It was fully connected to a two-dimensional output vector that was then forwarded to a ReLu function.

**Fig 2.**
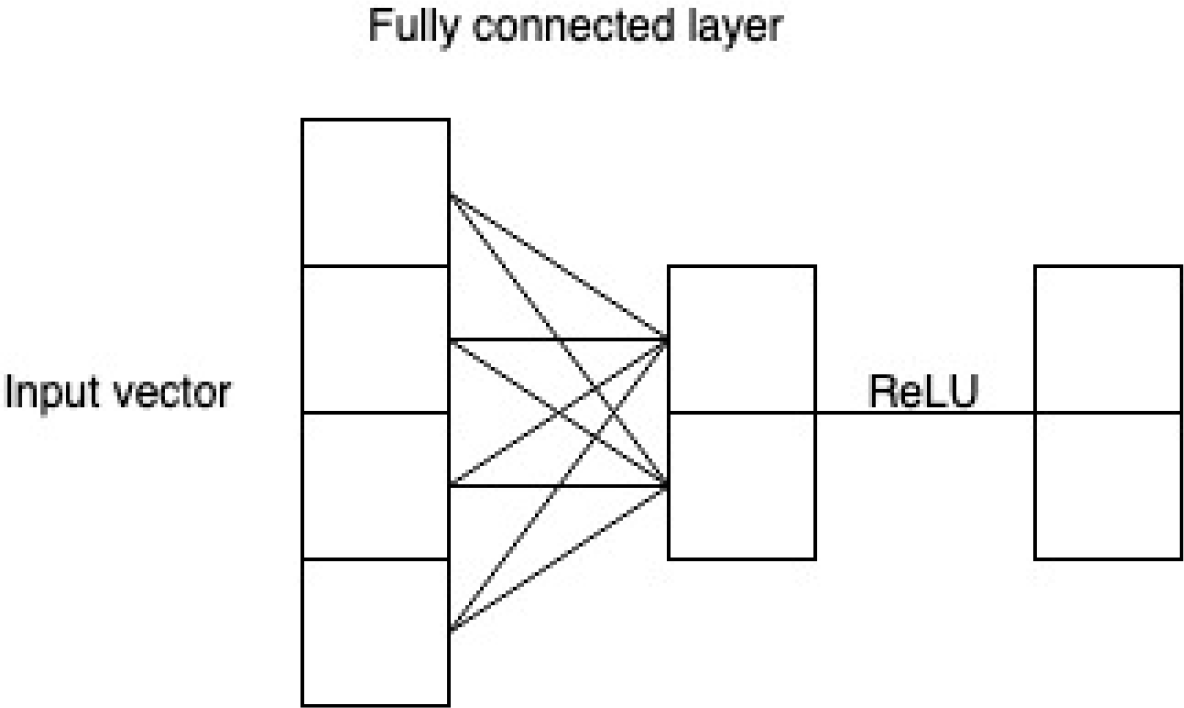
Multi-layer perceptron for attributes vector with rectified linear unit (ReLU) activation function. From left to right the MLP consists of a four-dimensional attributes vector, a two-dimensional vector, and a two-dimensional output vector.

#### Regression head

The mandatory regression head, which used temporal data as input, was constructed with five different neural network types: conventional recurrent neural networks (RNNs), convolutional neural networks, residual network, DenseNet, and Xception network. The output of every network type in the regression branch was a vector of size 16 or 32 that was used for regression or concatenation with the feature head.

#### Recurrent neural networks

For the conventional recurrent neural networks, recurrent neural network [16], long short-term memory [17], and gated recurrent units (GRUs; [18] layers were tested. These layers are the conventional approaches for problems using time-series data as input and were therefore used as benchmark. Each RNN configuration had three layers with each hidden layer containing 128 neurons. In the experiments presented in the next section both directional and bidirectional versions of the three layer types were tested.

#### Convolutional neural networks

In the experiment, a fully convolutional neural network (CNN), mentioned in [20], was also evaluated. The network contained three 1D-convolutional layers and a fully connected layer at the end. The output is a vector of size 16 or 32. The architecture of the CNN is displayed in Fig 3. Kernel size was set to three and the number of filters in the three convolutional layers were *n*_*f*_ = {32, 64, 32}. Two CNN configurations were implemented, one with 16 and one with 32 neurons in the last fully connected layer.

**Fig 3.**
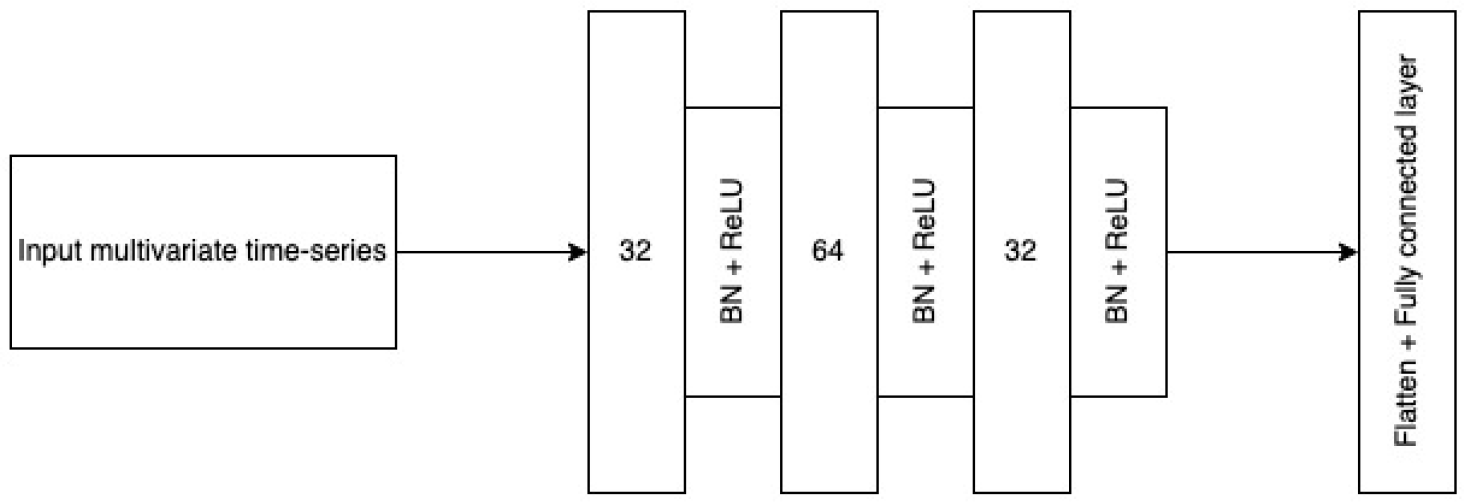
Architecture of the convolutional neural network. Batch normalization (BN) and ReLU activation function were used.

#### Residual neural networks

A residual network (ResNet) employs skip connections to facilitate the gradient flows during training. The architecture has been successfully applied to computer vision tasks [21], but also application on time series data have been studied (e.g. [20]). In this paper the architecture from [20] was used, with modifications to the hyperparameters to ensure that the model was compatible with the considerably smaller dataset.

The residual network was built from so-called residual blocks, which each contained three one-dimensional convolutional layers as well as a direct shortcut from input to output that used addition. In the experiment, three blocks with three convolutional layers each were used. For the kernel sizes *K*_ks_ in each layer two different options were tested, *K*_ks_ = {3, 3, 3} and *K*_ks_ = {7, 5, 3}. The number of filters in the three residual blocks were set to {*n*_*f*_, 2*n*_*f*_, 2*n*_*f*_} with *n*_*f*_ = 24. The dimension of the output vector (*c*_out_) was set to 16. The third block was followed by a global average pooling layer and a softmax layer. Fig 4 illustrates the tested network architecture.

**Fig 4.**
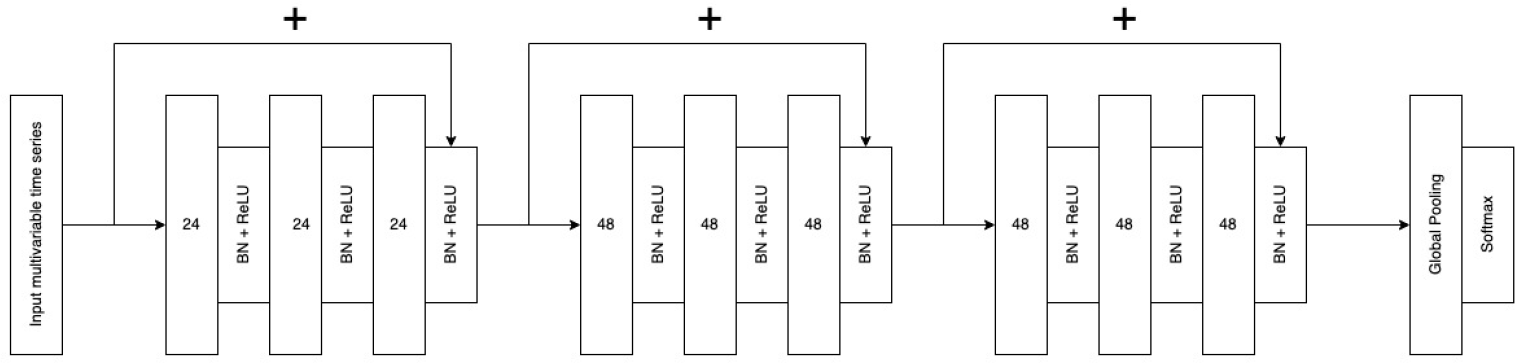
Architecture of the residual neural network, based on the architecture from [20].

#### DenseNet

The DenseNet architecture is inspired by the skip connections in ResNet. However, instead of summing input and output at specific shortcut as in ResNet, the DenseNet architecture uses dense connectivity: the feature map outputs of any layer are directly concatenate to subsequent layers [14]. Overall, the DenseNet architecture is divided into three levels from simple to sophisticated: densely connected layer (hereafter called dense layer), dense block, and dense architecture. The dense layer contains two convolutional layers. In this paper, one-dimensional (1D) layers are used for time series data instead of two-dimensional (2D) for computer vision as in the original paper [14]. The kernel sizes for convolutional layers were 1 and 3 respectively, and the number of filters was 16. The dense layer uses residual connection where output and input are added (see Fig 5(a)). Next, each dense block (see Fig 5(b)) is built from a positional encoding layer [22], four dense layers and a 1D max pooling layer. The dense layers are densely connected, which means the output of every previous layer is concatenated to the later layers. Finally, the DenseNet architecture (see Fig 5(c)) consists of a convolutional layer and four consecutive dense block. The convolutional layer transforms the time series input, a *l*-by-5 matrix with *l* being the length of the time series, into a feature map of size *l*-by-24 at the beginning of the network.

**Fig 5.**
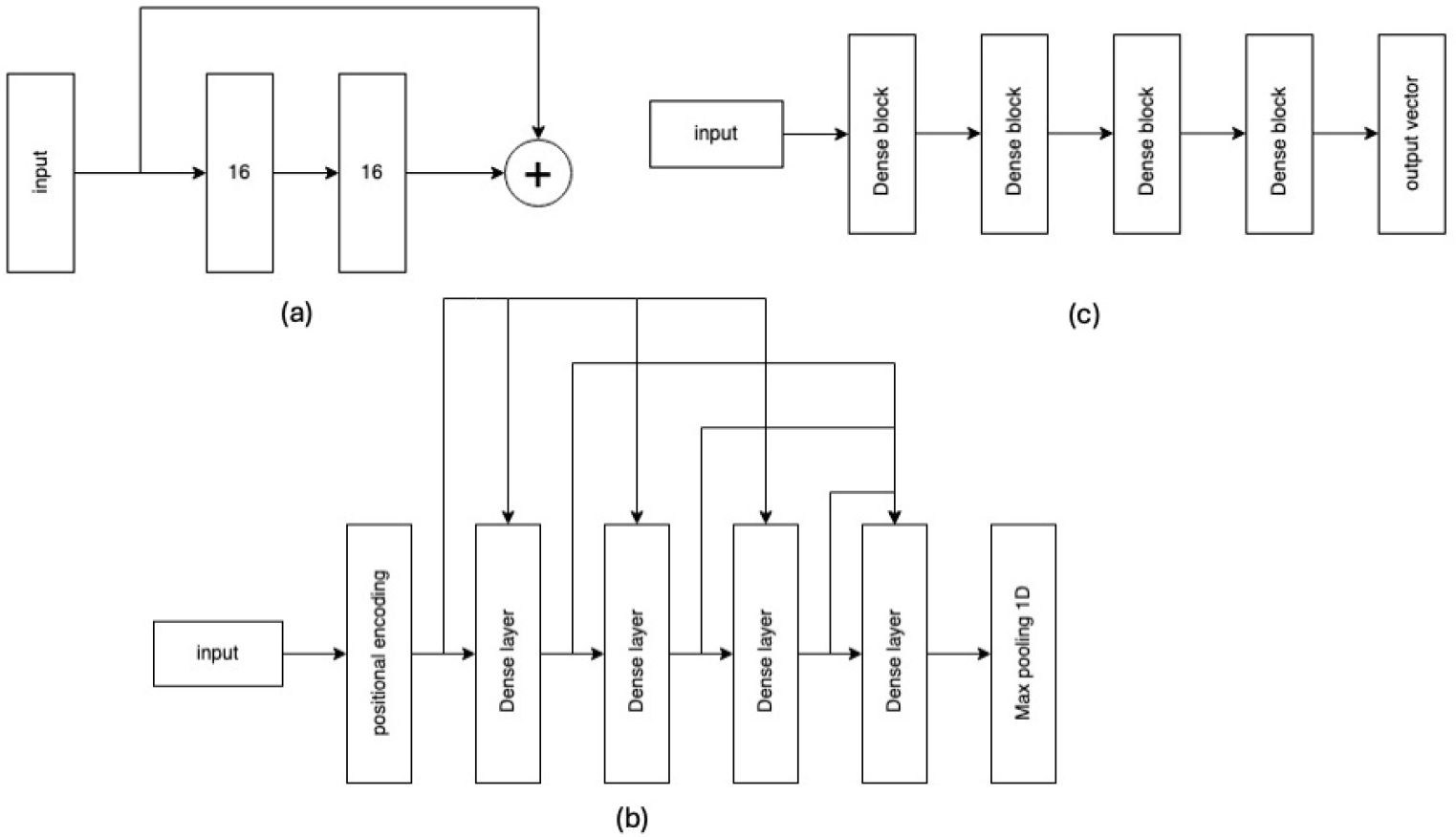
Architecture and building blocks of DenseNet. Dense layer architecture is shown in (a), dense block architecture in (b), and overall architecture in (c).

#### Xception network

The Xception network in this paper follows [23]. This network type consists of Xception modules that include two paths, a 3-depthwise separable convolution and a max pooling path, and are linked by residual connections. This architecture enables Xception networks to learn the input with various kernel sizes to obtain both the long and short-term structure in the data [23].

Fig 6(a) and Fig6(b) illustrate an Xception module and the Xception network architecture. In Fig 6(a) *f* @*SeparableConv*(*s, k, p*) denotes a depthwise separable convolution with *f* filters, stride *s*, which refers to the number of time steps the filter moves ahead in the input time series data matrix, kernel size *k*, and padding *p*, which refers to the process of adding artificial values outside the *l*-by-5 input data matrix to also cover the so-called border areas. Similarly, *f* @*Conv*1*x*1 denotes a convolution layer with *f* filters, stride 1, kernel size 1, and padding 0.

**Fig 6.**
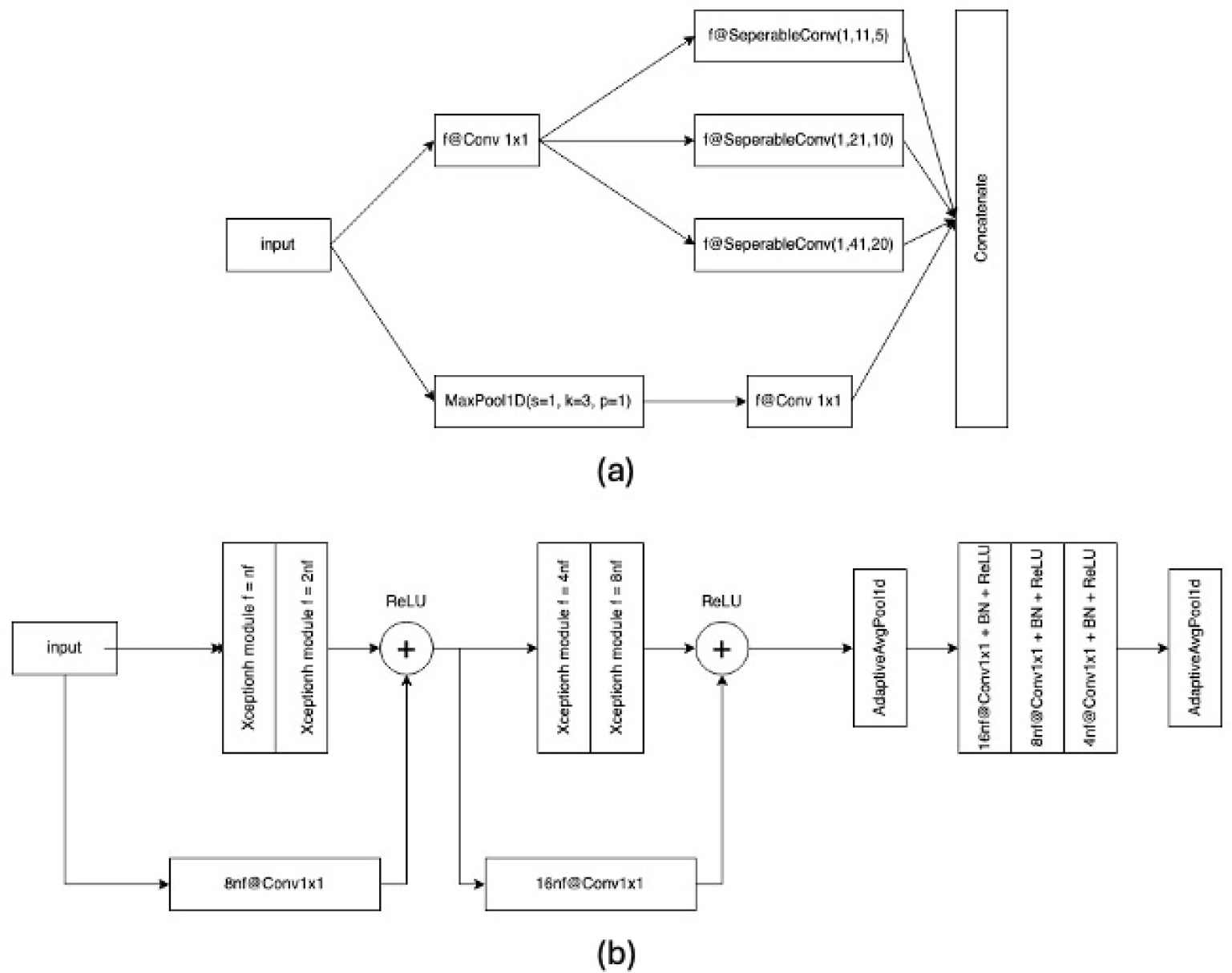
Architecture and building blocks of XceptionNet. Architecture of an Xception module is shown in (a), and the Xception network architecture architecture is illustrated in (b).

## Results and discussion

### Correlation analysis

The analysis in [8] revealed that the target feature oxygen consumption (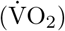) had highest correlations to input features speed, heart rate and speed change, while it was only weakly negatively correlated to vertical oscillation and step duration. Still, adding the latter two as input features to the *LSTM model* in [8] improved the accuracy of 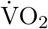 estimations somewhat.

One shortcoming of the analysis in [8] was that only Pearson correlation coefficients were computed, which indicate the strength of a linear relationship between two features but does not provide information on potential nonlinear statistical relationships.

Therefore, for this paper also Spearman correlation coefficients were calculated, in addition to Pearson correlation coefficients. Spearman correlation coefficients describe how well a monotonic function describes the relationship between two features.

The correlation coefficients for raw data are displayed in Fig 7. The Pearson correlation plot (Fig 7(a)) shows that heart rate, speed, and speed change have the highest linear correlation with oxygen consumption. It is interesting to note that the correlation coefficients for speed, vertical oscillation and step duration are approximately the same as in [8] (differences at most 0.05) but that the coefficients for heart rate and speed change were approximately 0.2 and 0.3 higher than in [8].

**Fig 7.**
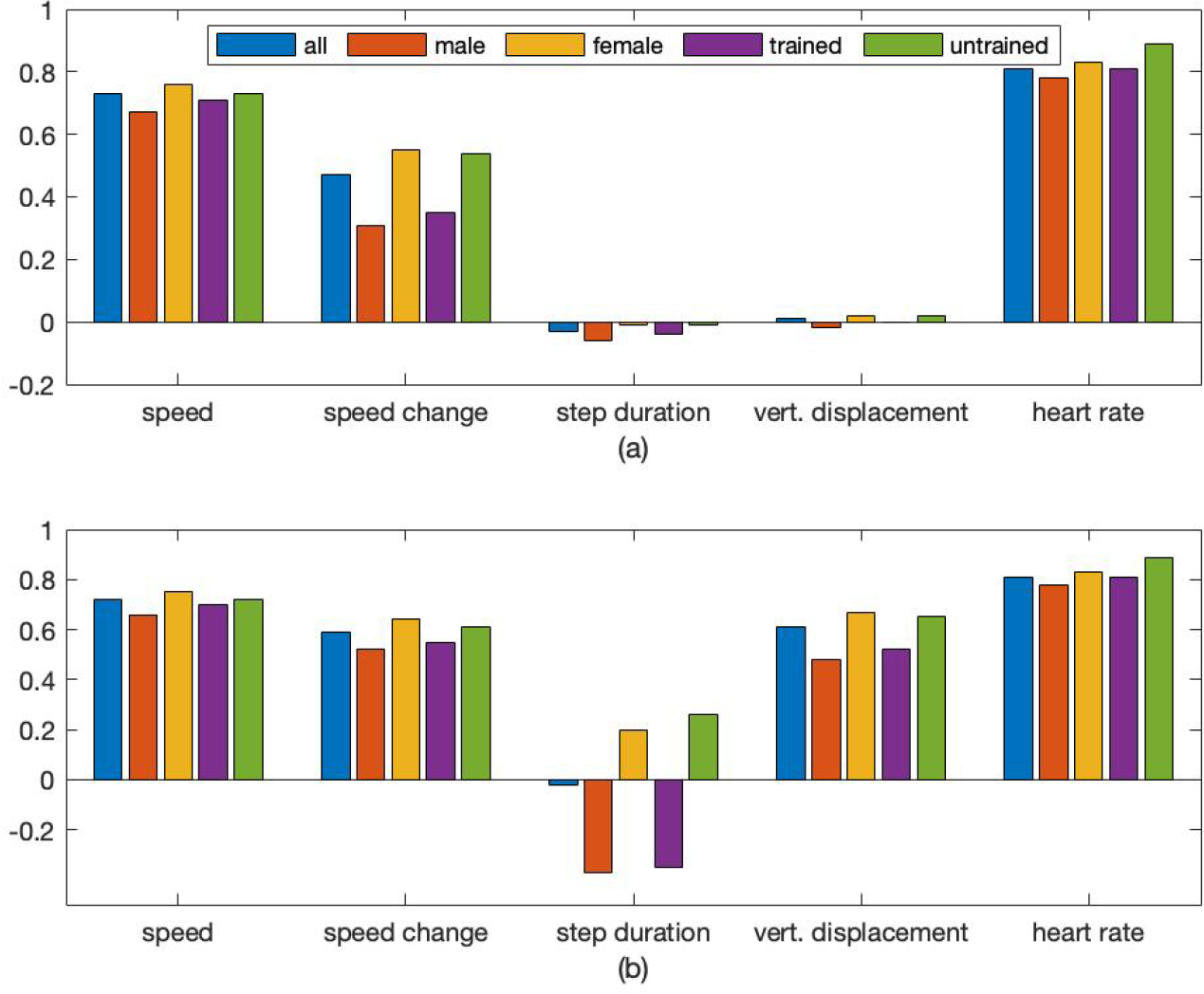
Correlation coefficients between oxygen consumption (target feature) and five input features. Input features included speed, speed change, step duration, vertical displacement, and heart rate. Pearson and Spearman correlation coefficients are shown in (a) and (b) respectively.

Looking at the Spearman correlation coefficients (Fig 7(b)) it can be noted that correlations between oxygen consumption and heart rate and speed respectively are (almost) the same as the Pearson coefficients, which could be expected due to a linear function being a monotonic function. For speed change the Spearman coefficient is 0.12 larger, supporting the assumption that the relationship between speed change and oxygen consumption is only approximately linear. The most interesting finding is, however, that vertical oscillation and oxygen consumption are strongly correlated (0.61). Together with the low Pearson coefficient (0.01) this indicates that the relationship between these two features is highly nonlinear and explains why adding it as predictor to the *LSTM model* in [8] improved the estimation accuracy. On the contrary, the Spearman coefficient of step duration and oxygen consumption is, similarly to the Pearson coefficient, approximately zero.

However, comparing the Pearson and Spearman correlations for data from only male with those for data from only female participants reveals a potential explanation why step duration is a useful input feature for estimating oxygen consumption. It is negatively correlated to oxygen consumption for male (−0.37) but positively correlated for female (0.2). This indicates that using step duration as an input feature could improve accuracy if gender is taken into account. Since the analysed dataset was gender balanced this relationship was hidden when considering the whole dataset. Other features that showed noticeable differences in correlation with oxygen consumption between male and female participants were speed change (both Pearson and Spearman coefficients) and vertical oscillation (only Spearman coefficient).

Interestingly enough, similar correlation values and differences as for male vs. female participants were found for trained vs. untrained participants (fitness level set to 1 vs. 0), even so both the trained and the untrained group had a male-to-female ratio of one. Furthermore, it was noticed that heart rate and oxygen consumption showed very high Pearson and Spearman coefficients for untrained participants, indicating an almost linear relationship between heart rate and oxygen consumption.

### Intra-subject estimations

For comparing the performance of the *Modified LSTM model* with the preformance of the *LSTM model* [8] the root mean square error (RMSE) for the 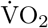 estimations over all 16 subjects was calculated. While the average RMSE for the *LSTM model* was 3.3459 ml×min^−1^×kg^−1^ (standard deviation: 2.3568 ml×min^−1^×kg^−1^), it was only 0.6019 ml×min^−1^×kg^−1^ (standard deviation: 0.3076 ml×min^−1^×kg^−1^) for the *Modified LSTM model*. Fig 8(a) and 8(b) show the Bland-Altman analysis for both model configurations. For the *LSTM model* the estimation bias was −0.4356 ml×min^−1^×kg^−1^, which is approximately 0.8712% of peak 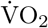, while the bias of the *Modified LSTM model* was only −0.0078 ml×min^−1^×kg^−1^, which is approximately 0.0156% of peak 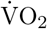. The validity of estimated oxygen consumption expressed by 95% limits of agreement wre 8.2292 ml×min^−1^×kg^−1^ (approximately 16.4584% of peak 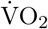) for *LSTM model* and 1.5470 ml×min^−1^×kg^−1^ (approximately 3.0939% of peak 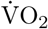) for *Modified LSTM model*. This shows that by some simple improvements to the architecture of the LSTM model its accuracy for intra-subject 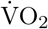 estimation can be improved considerably.

**Fig 8.**
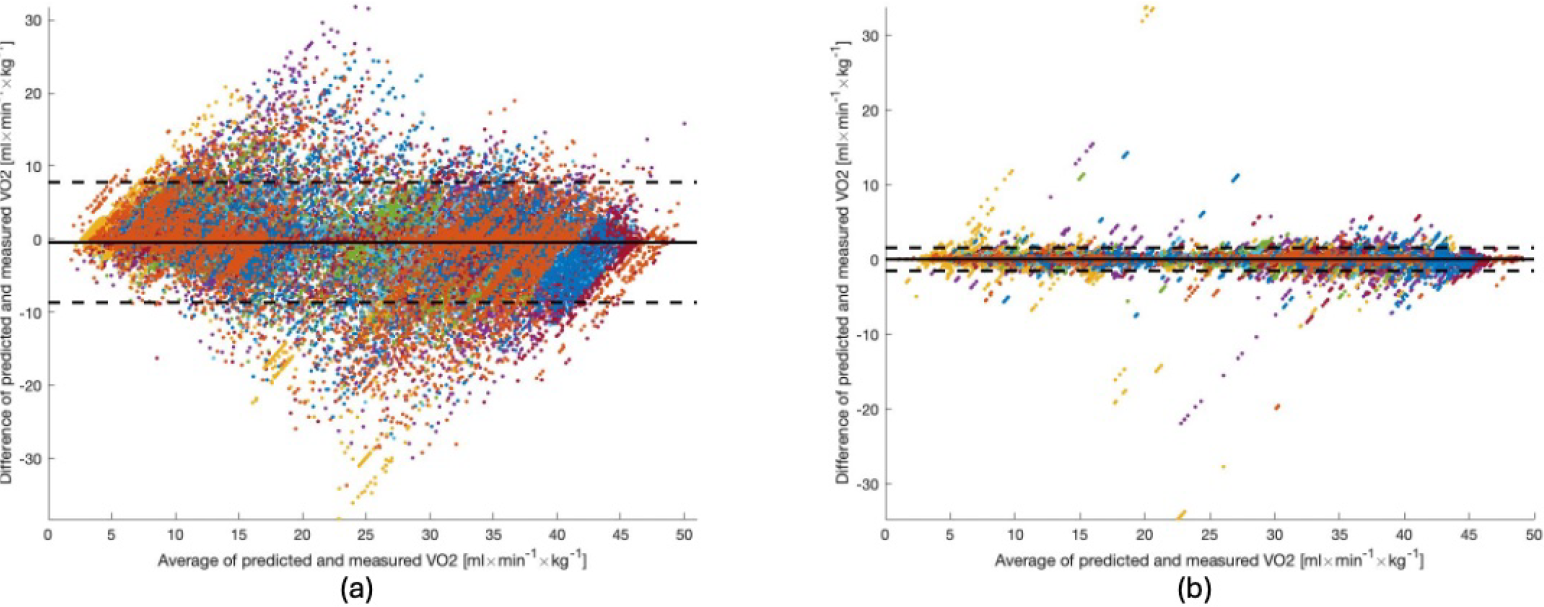
Bland–Altman analysis of the estimated and directly measured oxygen consumption. Subfigure (a) shows results for the *LSTM model* from [8] and subfigure (b) shows results for the *Modified LSTM* model with data from all 16 subjects. Dashed horizontal lines represent the 95% limits of agreement and solid lines represent estimation biases. Each color represents data from a unique participant in the test set. For better comparability the y-axes are equally scaled.

### Inter-subject estimations

Table 1 lists the results of a selection of network configurations for inter-subject oxygen consumption estimation, sorted with respect to the average root-mean square error of the oxygen consumption estimations over all cross-validation rounds (column *RMSE (mean)*).

**Table 1.**
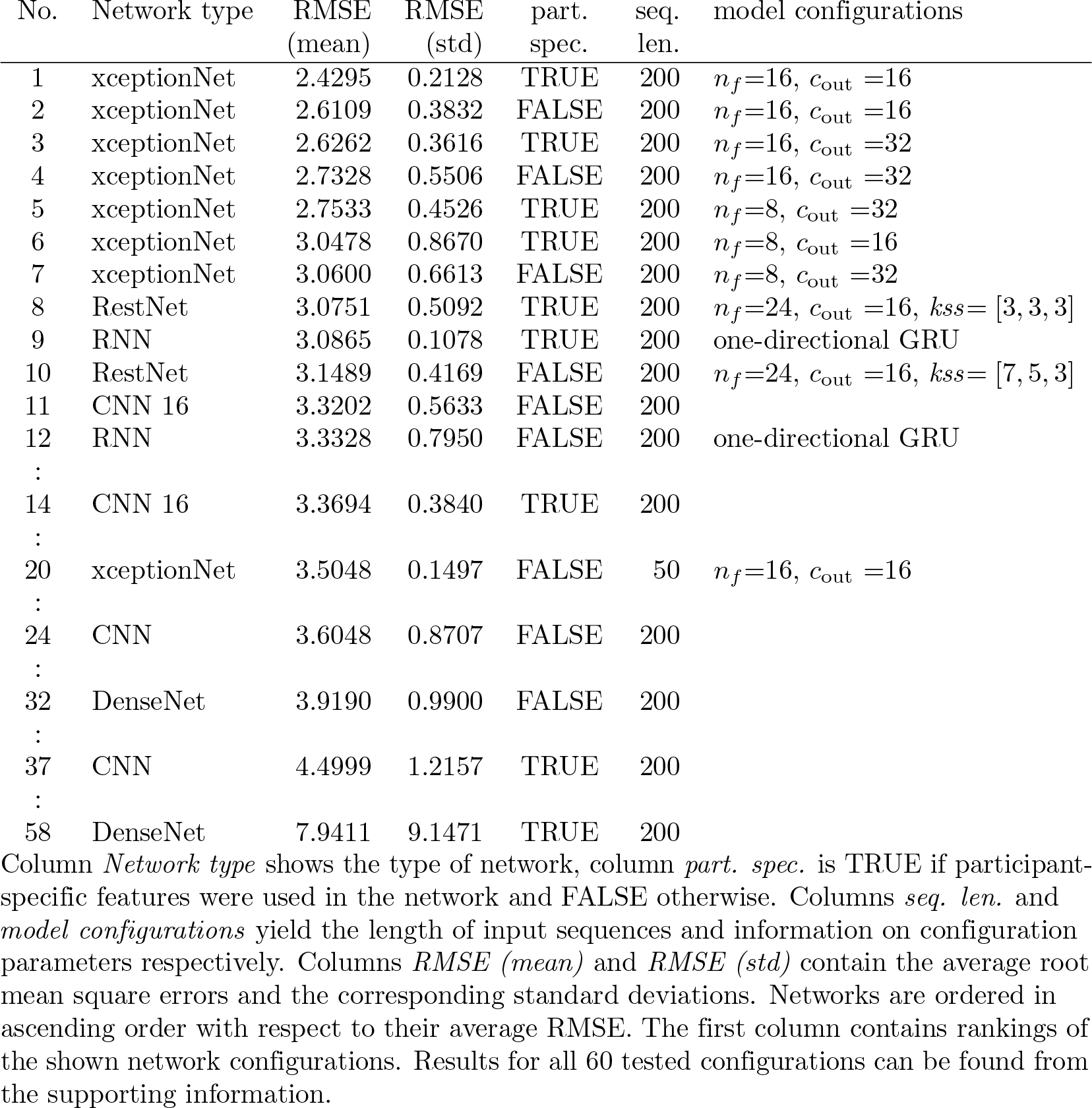
Results for a selection of neural network configurations for inter-subject estimation of oxygen consumption.

**Table 2.** Results for best 40 tested neural network configurations for inter-subject estimation of oxygen consumption.

| No. | Network type | RMSE<br>(mean) | RMSE<br>(std) | part.<br>spec. | seq.<br>len. | model configurations |
| --- | --- | --- | --- | --- | --- | --- |
| 1 | xceptionNet | 2.4295 | 0.2128 | TRUE | 200 | $n_f=16, c_{out}=16$ |
| 2 | xceptionNet | 2.6109 | 0.3832 | FALSE | 200 | $n_f=16, c_{out}=16$ |
| 3 | xceptionNet | 2.6262 | 0.3616 | TRUE | 200 | $n_f=16, c_{out}=32$ |
| 4 | xceptionNet | 2.7328 | 0.5506 | FALSE | 200 | $n_f=16, c_{out}=32$ |
| 5 | xceptionNet | 2.7533 | 0.4526 | TRUE | 200 | $n_f=8, c_{out}=32$ |
| 6 | xceptionNet | 3.0478 | 0.8670 | TRUE | 200 | $n_f=8, c_{out}=16$ |
| 7 | xceptionNet | 3.0600 | 0.6613 | FALSE | 200 | $n_f=8, c_{out}=32$ |
| 8 | RestNet | 3.0751 | 0.5092 | TRUE | 200 | $n_f=24, c_{out}=16, kss=[3, 3, 3]$ |
| 9 | RNN | 3.0865 | 0.1078 | TRUE | 200 | one-directional GRU |
| 10 | RestNet | 3.1489 | 0.4169 | FALSE | 200 | $n_f=24, c_{out}=16, kss=[7, 5, 3]$ |
| 11 | CNN 16 | 3.3202 | 0.5633 | FALSE | 200 |  |
| 12 | RNN | 3.3328 | 0.7950 | FALSE | 200 | one-directional GRU |
| 13 | RestNet | 3.3496 | 0.7900 | TRUE | 200 | $n_f=24, c_{out}=16, kss=[7, 5, 3]$ |
| 14 | CNN 16 | 3.3694 | 0.3840 | TRUE | 200 |  |
| 15 | RNN | 3.3844 | 0.4231 | TRUE | 200 | bi-directional RNN |
| 16 | RNN | 3.4680 | 0.6242 | FALSE | 200 | bi-directional GRU |
| 17 | RNN | 3.4716 | 0.4429 | FALSE | 200 | bi-directional RNN |
| 18 | RNN | 3.4911 | 0.4315 | FALSE | 200 | one-directional LSTM |
| 19 | RNN | 3.5010 | 0.7129 | FALSE | 200 | bi-directional LSTM |
| 20 | xceptionNet | 3.5048 | 0.1497 | FALSE | 50 | $n_f=16, c_{out}=16$ |
| 21 | RestNet | 3.5128 | 0.5988 | FALSE | 200 | $n_f=24, c_{out}=16, kss=[3, 3, 3]$ |
| 22 | RestNet | 3.5486 | 0.6032 | TRUE | 50 | $n_f=24, c_{out}=16, kss=[3, 3, 3]$ |
| 23 | RNN | 3.5719 | 0.6276 | TRUE | 200 | one-directional LSTM |
| 24 | CNN | 3.6048 | 0.8707 | FALSE | 200 |  |
| 25 | RNN | 3.6962 | 0.5952 | TRUE | 200 | bi-directional LSTM |
| 26 | RestNet | 3.7157 | 0.2107 | FALSE | 50 | $n_f=24, c_{out}=16, kss=[3, 3, 3]$ |
| 27 | RNN | 3.7365 | 0.4748 | FALSE | 200 | one-directional RNN |
| 28 | RNN | 3.7460 | 0.7335 | TRUE | 200 | bi-directional GRU |
| 29 | RestNet | 3.8792 | 0.7209 | FALSE | 50 | $n_f=24, c_{out}=16, kss=[7, 5, 3]$ |
| 30 | RestNet | 3.8856 | 0.8442 | TRUE | 50 | $n_f=24, c_{out}=16, kss=[7, 5, 3]$ |
| 31 | RNN | 3.9168 | 0.5130 | TRUE | 200 | one-directional RNN |
| 32 | DenseNet | 3.9190 | 0.9900 | FALSE | 200 |  |
| 33 | xceptionNet | 4.0829 | 0.7036 | TRUE | 50 | $n_f=16, c_{out}=32$ |
| 34 | xceptionNet | 4.0942 | 0.5790 | TRUE | 50 | $n_f=8, c_{out}=16$ |
| 35 | xceptionNet | 4.1529 | 0.6883 | FALSE | 50 | $n_f=8, c_{out}=32$ |
| 36 | xceptionNet | 4.3845 | 1.5622 | FALSE | 200 | $n_f=8, c_{out}=16$ |
| 37 | CNN | 4.4999 | 1.2157 | TRUE | 200 |  |
| 38 | xceptionNet | 4.6494 | 0.9080 | FALSE | 50 | $n_f=8, c_{out}=16$ |
| 39 | xceptionNet | 4.8867 | 0.6928 | TRUE | 50 | $n_f=16, c_{out}=16$ |
| 40 | DenseNet | 4.9339 | 0.6294 | TRUE | 50 |  |

Column *Network type* shows the type of network, column *part. spec.* is TRUE if participant-specific features were used in the network and FALSE otherwise. Columns *seq. len.* and *model configurations* yield the length of input sequences and information on configuration parameters respectively. Columns *RMSE (mean)* and *RMSE (std)* contain the average root mean square errors and the corresponding standard deviations. Networks are ordered in ascending order with respect to their average RMSE. The first column contains rankings of the shown network configurations.

**Table 3.** Results for 20 worst tested neural network configurations for inter-subject estimation of oxygen consumption.

| No. | Network type | RMSE<br>(mean) | RMSE<br>(std) | part.<br>spec. | seq.<br>len. | model configurations |
| --- | --- | --- | --- | --- | --- | --- |
| 41 | xceptionNet | 5.0884 | 1.3449 | FALSE | 50 | $n_f=16, c_{out}=32$ |
| 42 | CNN | 5.1862 | 0.3495 | TRUE | 50 |  |
| 43 | RNN | 5.2494 | 0.4595 | FALSE | 50 | bi-directional GRU |
| 44 | RNN | 5.2790 | 0.8909 | FALSE | 50 | bi-directional RNN |
| 45 | RNN | 5.3120 | 0.5529 | FALSE | 50 | one-directional GRU |
| 46 | RNN | 5.3302 | 0.8224 | FALSE | 50 | one-directional LSTM |
| 47 | RNN | 5.3399 | 0.7750 | FALSE | 50 | one-directional RNN |
| 48 | RNN | 5.4531 | 1.2189 | TRUE | 50 | bi-directional RNN |
| 49 | RNN | 5.4962 | 0.7417 | FALSE | 50 | bi-directional LSTM |
| 50 | RNN | 5.6207 | 1.0542 | TRUE | 50 | bi-directional GRU |
| 51 | RNN | 5.6606 | 1.3026 | TRUE | 50 | one-directional GRU |
| 52 | xceptionNet | 5.7136 | 2.3692 | TRUE | 50 | $n_f=8, c_{out}=32$ |
| 53 | RNN | 5.7538 | 1.1726 | TRUE | 50 | one-directional LSTM |
| 54 | CNN 16 | 5.7692 | 1.3642 | FALSE | 50 |  |
| 55 | RNN | 6.1492 | 2.0787 | TRUE | 50 | one-directional RNN |
| 56 | RNN | 6.3716 | 2.0295 | TRUE | 50 | bi-directional LSTM |
| 57 | CNN 16 | 6.5172 | 1.9349 | TRUE | 50 |  |
| 58 | DenseNet | 7.9411 | 9.1471 | TRUE | 200 |  |
| 59 | CNN | 8.1928 | 3.8141 | FALSE | 50 |  |
| 60 | DenseNet | 12.0831 | 8.9866 | FALSE | 50 |  |

Column *Network type* shows the type of network, column *part. spec.* is TRUE if participant-specific features were used in the network and FALSE otherwise. Columns *seq. len.* and *model configurations* yield the length of input sequences and information on configuration parameters respectively. Columns *RMSE (mean)* and *RMSE (std)* contain the average root mean square errors and the corresponding standard deviations. Networks are ordered in ascending order with respect to their average RMSE. The first column contains rankings of the shown network configurations.

For all configurations the average RMSE of test data and the corresponding standard deviations are shown. All five neural network types were tested with input sequences of 50 and 200 steps (column *seq. length*). In addition, the impact of using participant-specific features such as age, sex, body mass index and fitness level by enabling the optional MLP network for categorical variables was tested (variable *part. spec*. set to TRUE; if FALSE then participant-specific features were not used).

The most accurate 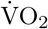 estimations were achieved by xceptionNet using sequences of 200 steps, participant-specific input features, *n*_*f*_ = 16, and 16 neurons in the output vector, with a RMSE of 2.4295 ml×min^−1^×kg^−1^ (standard deviation of 0.2128 ml×min^−1^×kg^−1^), which is better than the *LSTM model* from [8] used for intra-subject estimations in the previous subsection. Even without the use of participant-specific features the RMSE of this xceptionNet configuration increases only to 2.6109 ml×min^−1^×kg^−1^ (+7.47%; standard deviation of 0.3832 ml×min^−1^×kg^−1^). Even more promising is the fact that also other choices for *nf* and *c out* do not result in considerably worse performances. The seven best configurations are all xceptionNet configurations, which suggest that xceptionNet is the most promising network type for inter-subject oxygen consumption estimation.

Only in eight and ninth place the first non-xceptionNet configurations, a ResNet and a RNN configuration can be found. Their *RMSE test* are 26.58% 27.05% larger than that of the best xceptionNet. For CNN, the best accuracy was achieved with 16 neurons in the last layer before the regression layer, input sequences of 200 steps, and without using participant-specific features (11th best configuration). The best CNN with 32 neurons in the last layer before the regression layer (24th best configuration) yielded a 8.57% higher *RMSE test* than the best CNN 16. The worst network type for 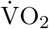 estimation is, based on this study, DenseNet, with the best configuration only yielding the 32nd best *RMSE (mean)* for the test data.

When it comes to length of input sequences, 200 steps seems to be a better choice than 50 steps, which is in contrast to the results in [8]. The best network configuration using input sequences of 50 steps is an xceptionNet with a *RMSE test* of 3.5048 ml×min^−1^×kg^−1^, which is 44.26% larger than the best overall network. The reason for the discrepancy between results from [8] and this paper is most likely that the data available for training was significantly smaller in [8], which only considered intra-subject 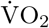 estimation.

Based on the study, no clear conclusion on the use of participant-specific features can be drawn. For example, for xceptionNet configurations omitting these features but using otherwise the same configuration resulted in six cases in 4.06% to 43.86% higher *RMSE test*, but in two cases the *RMSE test* was 37.58% to 39.43% higher when using participant-specific features than without them. For RNN configurations, however, using these features yielded in ten of twelve cases 2.32% to 15.93% higher *RMSE test* (for two cases it reduced the *RMSE test* by 2.51% to 7.39%).

## Conclusions

This paper had two aims. The first aim was to find techniques that would significantly increase the accuracy of intra-subject oxygen consumption estimations by LSTM network, compared to the LSTM network used in [8]. This earlier paper focused on demonstrating that LSTM networks can successfully estimate oxygen consumption, but refrained from optimizing the estimation performance. This first aim was achieved by including some early exit strategies into the LSTM network, reducing the number of hidden layers in the LSTM layer, and introducing an adaptive learning rate. Using these modifications, the average root-mean square error for oxygen consumption estimates was reduced by approximately 82% compared to the RMSE of the LSTM network from [8]. The corresponding standard deviation was reduced by approximately 87%.

These promising results encouraged us to investigate a more demanding task, namely developing neural networks that are able to provide accurate oxygen consumption estimates even for inter-subject estimations. Preliminary tries with the simple network structures from the first task yielded poor accuracy levels. Thus, our research focused then on studying more sophisticated, state-of-the-art neural network structures. Five different network types were tested with various configurations. The results suggest that it is indeed possible to accurately estimate the oxygen consumption of an individual from motion and heart rate data by neural networks even when the data on which the network is trained was collected from other individuals, in other environmental conditions. This second result is of higher importance as it has more relevance for real-world applications. The algorithms can be employed in portable devices for real-time assessment of oxygen consumption during walking or running, removing the constraints associated with the use of spirometers. Further use cases of the network types include continuous assessment and estimation of energy expenditure and aerobic fitness, continuous monitoring for early detection of fatigue and deterioration of physical health.

One shortcoming of the training procedure was that each participant walked/ran at the same speeds, although the anaerobic threshold (AT) speed can differ significantly. For example, in [25] the average velocity at AT (vAT) varied from 4.17 m/s to 5.36 m/s for male and from 4.17 m/s to 4.81 m/s for female elite distance runners. Therefore, it is reasonable to assume a lower average vAT and larger variation in vAT for the participants studied in this paper as the group included participants that identified as recreational runners and participants that did not. Thus, if training data was collected from subjects with high vAT but 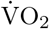 should be estimated for a subject with low vAT the estimation accuracy could suffer. Thus, training neural networks instead with data from vAT-based speeds, such as 25%, 50%, 75%,100% of an individual’s vAT might improve the estimation accuracy considerably. Testing this hypothesis is left for future research.

Future work should also focus on testing and, if necessary, adopting the estimation methods for very low and very high intensity exercises. Based on the correlation analysis, which revealed clear differences in correlations between step duration and 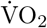 as well as speed change and 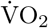 for male versus female, male/female specific models could be studied with larger datasets. In addition, the robustness of the estimation methods day-to-day variability in an individual’s oxygen consumption should be investigated. Finally, strategies for updating the estimation models with new unlabeled data should be developed. This last point has high relevance for real-world applications where for a new user in the beginning an estimation model will be used that had been trained with data from other individuals. However, once data from the new user, without reference 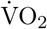 measurements, will be available the model could be fine-tuned to yield more accurate 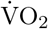 estimates in the future.

## Supporting information

**S1 Table. Results for best 40 tested neural network configurations for inter-subject estimation of oxygen consumption**.

**S2 Table. Results for 20 worst tested neural network configurations for inter-subject estimation of oxygen consumption**.

## Acknowledgments

This work was supported in part by the Academy of Finland, grants 287295 (under consortium “OpenKin: Sensor fusion for kinesiology research”) and 323472 (under consortium “GaitMaven: Machine learning for gait analysis and performance prediction”.

